# The time course of chromatic adaptation under immersive illumination

**DOI:** 10.1101/2020.03.10.984567

**Authors:** Gaurav Gupta, Naomi Gross, Ruben Pastilha, Anya Hurlbert

**Affiliations:** Institute of Neuroscience, University of Newcastle, Framlington Place, Newcastle upon Tyne, NE2 4HH, UK

**Keywords:** chromatic adaptation, achromatic adjustment, perceptual whitepoint, colour constancy

## Abstract

Chromatic adaptation is a major contributory mechanism to constancy, yet its extent depends on many factors - spectral, spatial and temporal - which vary between studies and hence may contribute to differences in reported constancy indices. Here, we use the achromatic adjustment method to characterise the temporal progression of chromatic adaptation under a wide range of illuminations in an immersive environment. We control both the spectral properties of the illumination at the eye and the spatial context of the adjusted surface, to disentangle global adaptation from local contrast effects. We measure the timecourse of chromatic adaptation by assessing achromatic adjustments in 6 discrete time slots over 340 seconds. We find that the change over time of the adaptation state, proximally indicated by colour constancy indices (quantified by the relative closeness of the perceptual whitepoint to the test illumination chromaticity), (a) can be modelled by a proportional rate growth function, typically requiring more than 5 minutes to stabilise; (b) depends on the contrast between the test surface and its background, specifically increasing with decreasing test-background contrast; and (c) is generally similar in both extent and rate for different test illumination chromaticities. Adaptation progression does not differ significantly between illuminations on or off the daylight locus. Our results highlight the importance of considering exposure duration and stimulus configuration, as well as the distance between the pre-adaptation (reference) and test illumination chromaticities, when using achromatic adjustment as a measure of colour constancy.

## 1. Introduction

Chromatic adaptation is a fundamental driver of colour constancy, the perceptual stability of object colours under varying illumination. However, multiple neural mechanisms are thought to underpin colour constancy [1, 2], acting at different sites and levels of visual processing, from adaptation of the retinal photoreceptors to extraction of second-order image information (such as specular highlights) to activation of object-specific memory modules in higher visual cortex [3]. It has previously been suggested that these mechanisms are optimised for natural environmental variations [4], and that constancy is better for illumination changes along the locus of daylight chromaticities (the “daylight locus”) than in other directions [5, 6]. Yet systematic studies of multiple types of illumination change are relatively rare, and their conclusions as well as methodologies differ [7]. In this study, we use the method of achromatic adjustment method to characterise the timecourse of chromatic adaptation under a wide range of illuminations, including simulated natural daylights, in a naturalistic immersive environment.

The achromatic setting, or perceptual whitepoint, is a widely used measure of colour constancy, representing the surface chromaticity that is contextually perceived as neutral. If chromatic adaptation and/or colour constancy were perfect, a spectrally non-selective surface, i.e. one that reflects 100% of the incident light at each visible wavelength, would continue to appear white, despite (or indeed due to) perfectly reflecting the ambient illumination, whose power may vary considerably across the spectrum. The closeness of the perceptual whitepoint to the ambient illumination chromaticity is therefore taken to be a measure of the non-exclusive phenomena of chromatic adaptation and colour constancy.

Chromatic adaptation occurs under light that differentially activates the three distinct cone types, and is typically modelled as a change in their relative response gains. The gain changes are typically derived from the perceptual whitepoint. For example, the “equivalent illumination” model of colour constancy [8] assumes that the effect of an illumination change is captured by a linear transformation of the three cone excitations, in which the gains are derived from the relationship between the observer’s perceptual whitepoints under the reference and test illuminations. Similarly, standard colour appearance models [9, 10] incorporate a chromatic adaptation transform which converts the tristimulus coordinates (themselves linear transformations of the receptoral responses) of a surface under a test illumination into their equivalent under a reference illumination, normalising them by the tristimulus values for a white surface under the test illumination, and with a subsequent non-linear luminance adaptation transform. The underlying assumption is that the adaptation transform fully accounts for the effects of the test illumination chromaticity on colour appearance. Complete constancy is therefore synonymous with complete chromatic adaptation, when degree of adaptation D is 1.

These models in turn are related to von Kries adaptation, which also forms the basis for computational models of colour constancy that recover invariant surface descriptors via linear transformations of the sensor responses [11]. Adaptation of colour-selective neurons at more distal levels of the visual pathways also occurs [12–15] and therefore it is appropriate to consider chromatic adaptation more broadly as a perceptual phenomenon underpinned by physiological mechanisms at multiple levels in visual processing. Here we use the term chromatic adaptation in this sense and do not attempt to disentangle retinal from higher-level contributions, or to further develop chromatic adaptation transforms. Instead, we focus on the temporal progression of adaptation and the effect of ambient illumination and patch-surround chromaticities.

The extent of chromatic adaptation is known to depend on multiple factors: duration of adaptation [16–19], spatial and chromatic characteristics of the surround [20, 21], chromaticity and intensity of adapting light [22, 23], and intensity of the test stimulus [24]. Correspondingly, the extent of colour constancy as measured by the perceptual whitepoint also depends on these factors, and therefore differences in colour constancy indices reported from achromatic adjustment studies may also be due to variations in these factors [8,23,25]. In particular, reported differences in the chromatic bias of achromatic points and their dependence on the chromaticity or intensity of the adapting light [8, 22, 23] may be partly explained by crucial differences in adaptation duration and surround conditions, as well as by differences in viewing conditions.

Studies agree that chromatic adaptation occurs on distinct time scales, from seconds and minutes (“short-term”) to hours and days (“long-” and “very-long-term”) [19], supporting the hypothesis that multi-level neural mechanisms contribute to adaptation. Short term adaptation itself consists of distinct slow and fast response phases [16, 17]. Fairchild (1995) [16] find chromatic adaptation at constant luminance to be 90% complete after approximately 60s, with a sum of exponentials fit to the data. Rinner and Gegenfurtner (2000) [17] find a half-life of approximately 20 seconds for the slow phase of short-term adaptation, with changes in the perceptual whitepoint continuing up to the longest duration tested of 120 seconds, based on a single exponential model fit. In these experiments, the change from reference to adapting illumination was always from one extreme of a cardinal cone-opponent axis to the other (i.e. red-green or blue-yellow), and there was no difference in adaptation time-course between cardinal directions. Rinner and Gegenfurtner (2000) [17] also find two phases of adaptation: (1) a fast phase, with half-life of 67 ms, likely to occur via receptoral gain adjustments and (2) an effectively instantaneous phase, operating in the first 25 ms, accounting for 60% of the total adaptational effect on colour appearance, and attributed to cortical spatial contrast processes. Shevell (2001) [18] demonstrates that variations in the surround interact differentially with these distinct adaptation phases (see also Werner (2014) [21]), with the spatial context having greater influence on the slow phase, compared to the fast, up to a steady state at 5 minutes or beyond.

In these studies the final perceptual whitepoint was not expressly compared with the adapting illumination chromaticity in order to extract a direct measure of colour constancy. The completeness of constancy for different adapting lights is therefore indeterminate. Other studies, which directly derive colour constancy indices from perceptual whitepoints, report varying extents of chromatic biases in adaptation, particularly with respect to the hue blue. Brainard (1998) [8] measures achromatic adjustment of a test patch, controlled by a projection colourimeter and enclosed within a grey surround, after 20 seconds of adaptation to each of 11 illuminations, and with no reported restrictions on the adjustment period. A high level of adaptation completeness is observed, with similar levels of completeness for all illumination hues. In contrast, Weiss et al. (2017) [23] test chromatic adaptation to an on-screen surround for 40 simulated illumination hues, using untimed achromatic adjustment in a patch-surround setup, and report almost all achromatic adjustments being shifted towards blue. Lee et al. (2012) [26] measure the shift in the achromatic point with changing illumination via the intersection of blue-yellow and red-green judgements for 120 surfaces, two backgrounds (black and illuminated), two illuminations (sunlight yellow and skylight blue), over three observers and a timecourse of 21 seconds per illumination. They find a slower update of the achromatic point for the black background, a temporal influence of patch chromaticity, and generally incomplete adaptation over 21 seconds.

Previous studies also differ in their conclusions on the effects of the test stimulus intensity. Kuriki (2006) [24] demonstrates that the perceptual whitepoint shifts away from the illumination chromaticity and towards equal energy white as the intensity of the test stimulus increases. The stimulus background being held at a constant luminance brighter than the darkest test stimulus, this result means that constancy decreases as the stimulus becomes brighter than its immediate surround. Similarly, Werner and Walraven (1982) [22] report that achromatic points shift more towards the adapting background chromaticity as the luminance of the background increases and contrast with the test stimulus decreases. Brainard (1998), however, finds no effect of background luminance on the achromatic point for a given test illumination chromaticity.

Although Kuiriki (2006) do not report colour constancy indices or distances of achromatic points from illumination chromaticity, their data indicate that even the settings at the lowest test patch intensities, which would display greatest constancy on this measure, are significantly different from the illumination chromaticity. These results differ from Brainard’s (1998) report of near-perfect constancy, as measured by the equivalent illumination chromaticities, for 8 illuminations off the daylight locus. Neither Brainard et al (1998) nor Kuriki (2006) constrained or recorded the duration of the achromatic adjustment periods so it is unclear whether the differences in constancy are due to different temporal extents of chromatic adaptation.

Surround effects may also contribute significantly to differences between results across all studies. Both the local and global surround influence colour appearance, with the greatest effect arising from the immediate vicinity of the focal patch and local edge contrasts, and a declining but significant effect for more remote surrounds [20, 27]. The size and content of the adapting field are therefore important factors, and these differ significantly between studies, from fully isolated small-field stimuli in Maxwellian view [22] to midsize-field computer monitor displays (e.g. ∼ 60° by 40° [17, 23]; ∼38° by 30° [26]), to fully immersive illumination environments [8, 24].

The sign and magnitude of local edge contrasts between test stimulus and surround are also critical, with some effects (e.g. the gamut expansion effect, [27]) being maximal for minimal contrast, and strongly reduced by the insertion of black borders between stimulus and surround [27, 28]. These effects, though, are confounded by the potential of the surround to supply a neutral reference surface from which the illumination chromaticity may simply be inferred [25]; for example, the stimulus configuration of Weiss et al. (2017) includes luminance-noise backgrounds, while Brainard’s (1998) configuration includes multiple neutral surfaces of differing luminances. Disentangling spatial contrast effects due to the surround from the potential contribution of neutral reference surfaces to higher-level inference therefore requires independent control of the spatial surround and the ambient illumination.

In this study, we measure the temporal progression of chromatic adaptation, by achromatic adjustment of a small test patch (an emissive surface masquerading as a material surface) in an immersive environment in which the illumination reaching the eye is constant across two different local surround conditions, one a neutral reference surface, and the other black. We use spectrally tuneable lamps to provide a range of test illuminations in order to compare the characteristics of the time course of chromatic adaptation for illumination chromaticities near and far from neutral, both on and off the daylight locus, and to determine adaptation levels via constancy indices for near-final adaptation points.

## 2. Methods

### 2.1. Overview

We measured the timecourse of chromatic adaptation under fourteen different ambient illuminations and for two surrounds (black and grey), by recording the change in participants’ perceptual whitepoints following the onset of each illumination. Participants were asked to adjust a displayed surface to neutral using free navigation via a gaming controller, while being immersed in chromatic ambient illuminations, for a block of six contiguous adaptation periods each followed by a time-limited adjustment slot, per illumination. Adaptation periods and adjustment slots were of fixed duration in order to obtain comparable adaptation measures over time between individuals and illuminations. Achromatic matches and adjustment times were recorded and analysed to establish the temporal progression and maximal extent of chromatic adaptation for each distinct illumination chromaticity. Achromatic adjustment results were collected for both black and grey surrounds, to explore the effect of the surround on the adaptation process. The ambient light illuminance and device screen luminance were held constant such that the 10 cd /m^2^ screen luminance closely matched the luminance of the grey surround in its immediate vicinity, under all test illuminations of 100 lux illuminance at the eye for both surrounds. Participants adjusted only the chromaticity of the screen.

The experiments were conducted in a 2m by 2m by 2.09m room, the “lightroom”, with white walls and four overhead 12-channel LED lamps, the sole illumination source, spaced evenly over the central horizontal axis of the room. A table was placed flush against the back wall of the room, a height-adjustable tray placed on top of it, and both the table and the tray covered with a black cloth. A custom-built wooden frame consisting of a flat base and a vertical face at one end was placed on top of the black cloth. The vertical face of the frame consisted only of four edges, to the back of which another solid wooden board holding a tablet or smartphone could be clamped, and to the front of which forward-facing sheets, each with 5cm by 9cm central viewport cut-outs, could be attached. The thickness of the edges and relative positioning of the device provided 3.5cm distance between the device screen and the front-facing cut-out, the recessed position minimising incident ambient light on the screen. Strips of Velcro were glued to the front-face of each of the frame edges, and counterpart strips glued to the reverse faces of a black and a grey surround board laser cut from large sheets of cardboard. The black surround was further painted over with black paint. The black surround measured 58cm by 59.5cm (54° by 55°); the grey surround 46cm by 47cm (44° by 45°).

Participants’ viewing angle and distance were kept stable using a height-adjustable chinrest attached to a second table in front of the first, such that the viewing distance from the chinrest to the cut-out viewports on each surround was approximately 57cm, at which the rectangular patch subtended a visual angle of 5° by 9°. This table was also covered with black cloth. A fixed-height chair with static legs was placed in front of the chinrest, on which participants were instructed to sit and adjust the chinrest height to a comfortable viewing position. For each participant, the height-adjustable tray holding up the wooden frame was adjusted to match the viewing position such that the view from the participant’s eyes to the cut-out was horizontal.

Participants were draped with black cloth, from shoulders down, and were seated and positioned at the chinrest. On the table in front of them was placed an Xbox 360 gaming controller, using which they would make adjustments to the chromaticity of the display as visible through the viewport cut-out in the surround cardboard. From the chin-rest viewing position, the recessed display “patch” resembled a reflective flat surface, like paper. Although the participants were aware that it was an electronically controlled surface, the resemblance to a reflective surface was, by participant and experimenter account, remarkably strong. The participant’s task was to adjust the visible display patch to be a “neutral” in colour appearance under the test illuminations.

### 2.2. Participants

Eight participants (4 females and 4 males), with ages between 19 and 62 (mean age 31), were recruited from the student and general population of Newcastle upon Tyne, UK. Each was screened for normal or corrected visual acuity (by distance reading check; with usual corrective lenses as necessary) and normal trichromatic colour vision. Colour vision screening was performed using the Farnsworth Munsell 100 hue test, using lab-based age norms (consistent with published norms [29]) and the Ishihara 38 plates colour vision deficiency test, requiring fewer than two errors and none consistent with any colour vision deficiency.

Participants were required to attend two experiment sessions, each lasting two hours and scheduled on different days, with typically a few days to a week between sessions. Participants were randomly assigned the grey or black surround for their first session, and the other surround for the next. They received gift vouchers as compensation for their time.

### 2.3. Ambient illumination generation and calibration

The lightroom ambient illumination was generated by sending pre-calculated weights to each of the twelve independent channels of the LED light engine, which operates on the principle of pulse width modulation (PWM), with 1500 Hz controller frequency. Fractional weights in the range 0 to 1 modulate the channel outputs between 0 and 100% via PWM. The weights required to generate an illumination of arbitrary desired CIE tristimulus coordinates were calculated using the set of twelve spectral basis functions corresponding to the spectral outputs of the twelve channels at full power, 2° XYZ colour matching functions (based on the CIE 2006 2° LMS cone fundamentals [30]), and a spectral smoothness constraint [31] using Matlab’s quadratic programming function to optimise the solution.

Calibration measurements of the basis functions were carried out with an illuminance spectrophotometer (Konica Minolta CL500A) placed at the approximate mid-point between observer left and right eye positions. Prior to calibration, the LED lamps and CL500A were powered on (the lamps at experiment illuminance of 100 lux) and warmed up for at least thirty minutes. Calibration was performed separately for black and grey surrounds, ensuring the ability to achieve the same illuminance and chromaticity at the eye for each surround over all the test lights. Following calibration, we performed a calibration check routine that calculated the weights required for a gamut-encompassing discrete set of illuminations (3482 evenly-spaced Yxy grid points), and stepped through these, measuring the actual illumination with the CL500A, and computing *δE*_*uv*_ values between the target (intended) illuminations *T*_*i*_ and actual (measured) illuminations *M*_*i*_, with target illumination *T*_*i*_ as the adapting reference1, or 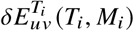. With the CL500A at the calibration position, and for the fixed at-eye illuminance level of the experiment, we obtain 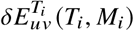 values less than ∼4, with a mean of ∼2, for both black and grey surrounds.

### 2.4. Device display colour calibration and adjustment

The device screen was calibrated by measuring channel gamma ramps using a computer-controlled spectroradiometer (Konica Minolta CS2000), and obtaining the mapping from RGB to CIE tristimulus coordinates using the 2° XYZ colour matching functions. A calibration check procedure iterated through a gamut-encompassing grid of discrete colorimetric gamut values (794 evenly-spaced Yxy grid points), displaying the equivalent RGB on the device screen, measuring the output with the CS2000, and computing the 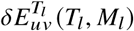 between intended *T*_*l*_ and measured *M*_*l*_ colorimetric values. Prior to calibration, the Tab S4 and CS2000 were powered on (the screen at experiment luminance of 10 cd /m^2^) and warmed up for at least thirty minutes. With the CS2000 at the calibration position2, and at the fixed display luminance level of the experiment, we obtain 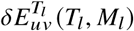 values less than ∼2, with a mean of ∼1, between predicted and measured values across the chromaticity gamut, allowing us to subsequently present patches of arbitrary chromaticity on the device display to high accuracy. The recessed position of the device screen minimised incident ambient illumination, and the calibration check routines performed under distinct test illuminations confirmed that there were no discernible effects of the ambient illumination on screen chromaticity.

The display device was set to keep the screen awake, with sleep disabled, and to operate at full power at all times. The device chosen for the experiment was the Samsung Galaxy Tab S4 running Android 8.1, for the quality of its sAMOLED display, 64GB memory and octacore processor. During a trial, all participant responses on the controller were recorded with timestamps. Navigation input via the controller were applied as stepped chromatic changes in CIELAB colour space, chosen for its near perceptual uniformity, along the *a* axis (red-green) for right and left controller inputs, and along the *b* axis (blue-yellow) for down and up controller inputs. The navigation step size was set small enough to produce colour changes of within one 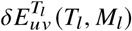 at the experiment luminance level. Participants were able to induce large changes in chromaticity by holding down direction keys on the controller, which resulted in a rapid sequence of small changes being registered, thereby allowing fast initial adjustment. For precision adjustments, participants were instructed to tap out the direction buttons, producing chromaticity changes of small step size. Participants’ final match submissions using the controller were blocked for three seconds after each submission, preventing accidental multiple submissions.

### 2.5. Test illuminations

A set of fourteen illuminations were selected, falling within the boundaries of the sub-gamut defined by shrinking the largest common gamut between the display device and the LED light engine to 0.75 of the distance from illuminant E or equal-energy white (hereafter refered to as EEW), at a common luminance level between the device screen and the immediately adjacent surface on the grey surround board. The test illuminations were selected from the reduced common gamut to allow sufficient navigation space, within the screen gamut, on all sides of the illumination chromaticity coordinates, thereby allowing the possibility of an exact screen chromaticity match to each test illumination chromaticity.

The set of test illuminations (Figure 1) was chosen to include extreme, moderate and neutral chromaticies, on and off the daylight locus. Extreme illuminations consisted of three “corner” chromaticities (lights 11, 10 and 12, approximately corresponding to red, green and blue respectively) and three mid-points along the sub-gamut edges (lights 14, 2 and 13, approximately corresponding to yellow, cyan and purple respectively). Moderate illuminations consisted of six chromaticities^3^ around D65 (lights 3 and 5-9), while the neutral illumination was D57 (light 4). Of the total set of illuminations, five fell on or sufficiently near the blackbody and daylight loci^4^ (lights 1, 7, 4, 5, 14, approximately corresponding to very blue, blueish, neutral, orangish and very orange/yellow). The total illuminance at the eye for each test light was 100 lux for both black and grey surrounds. The LED and device screen gamuts, with test illumination hues within the sub-gamut, are shown in Figure 1. Colorimetric specifications of the test illuminations, and chromatic differences (under experiment adaptation point D65) of test light *T*_*i*_ from EEW 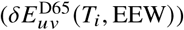 and from D65 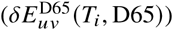, are given in Table 1. Illumination spectra are given in Figure 2.

**Table 1.** Test illumination colorimetry

| L | CIE x | CIE y | $\delta E_{uv}^{EEW}$ | $\delta E_{uv}^{D65}$ | CCT |
| --- | --- | --- | --- | --- | --- |
| 1 | 0.254 | 0.261 | 83 | 71 | 19774 |
| 2 | 0.270 | 0.380 | 76 | 62 | 8038 |
| 3 | 0.350 | 0.285 | 55 | 66 | 4313 |
| 4 | 0.328 | 0.344 | 12 | 13 | 5700 |
| 5 | 0.381 | 0.386 | 44 | 57 | 4053 |
| 6 | 0.281 | 0.339 | 49 | 33 | 8189 |
| 7 | 0.293 | 0.303 | 37 | 24 | 8221 |
| 8 | 0.385 | 0.339 | 46 | 64 | 3530 |
| 9 | 0.299 | 0.388 | 60 | 50 | 6713 |
| 10 | 0.333 | 0.536 | 125 | 121 | 5511 |
| 11 | 0.536 | 0.333 | 201 | 218 | 1762 <sup>a</sup> |
| 12 | 0.206 | 0.206 | 153 | 141 | $3 \times 10^5$ <sup>a</sup> |
| 13 | 0.401 | 0.266 | 120 | 133 | 2226 |
| 14 | 0.463 | 0.411 | 100 | 116 | 2652 |

<sup>a</sup>CCT approximated at these coordinates

**Fig. 1.**
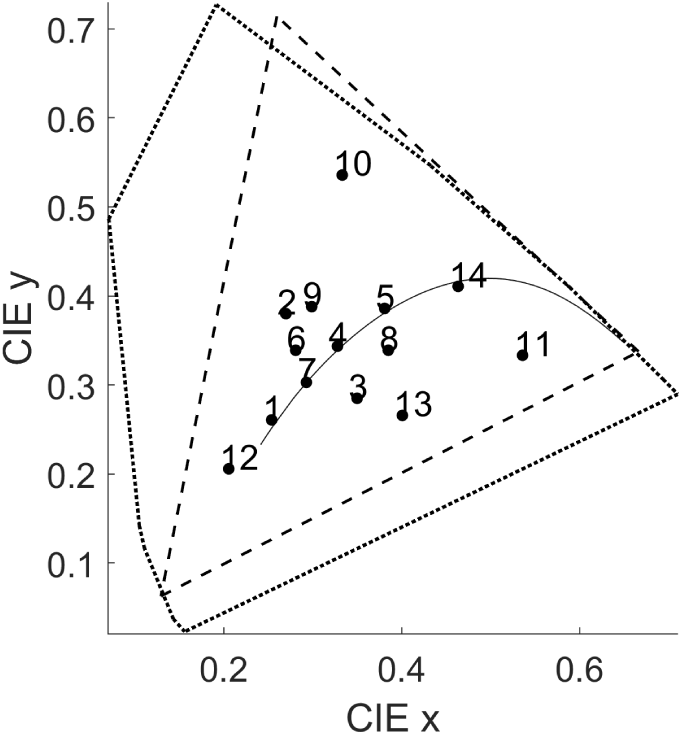
LED gamut (dotted polygon), screen gamut (dashed triangle) and illuminations. On daylight locus (black curve): 1, 7, 4 (D57), 5, 14. Moderate chromaticities: 7, 6, 9, 5, 8, 3. Extreme chromaticities: 10, 11, 12 (sub-gamut corners) and 2, 13, 14 (mid-points of corners).

**Fig. 2.**
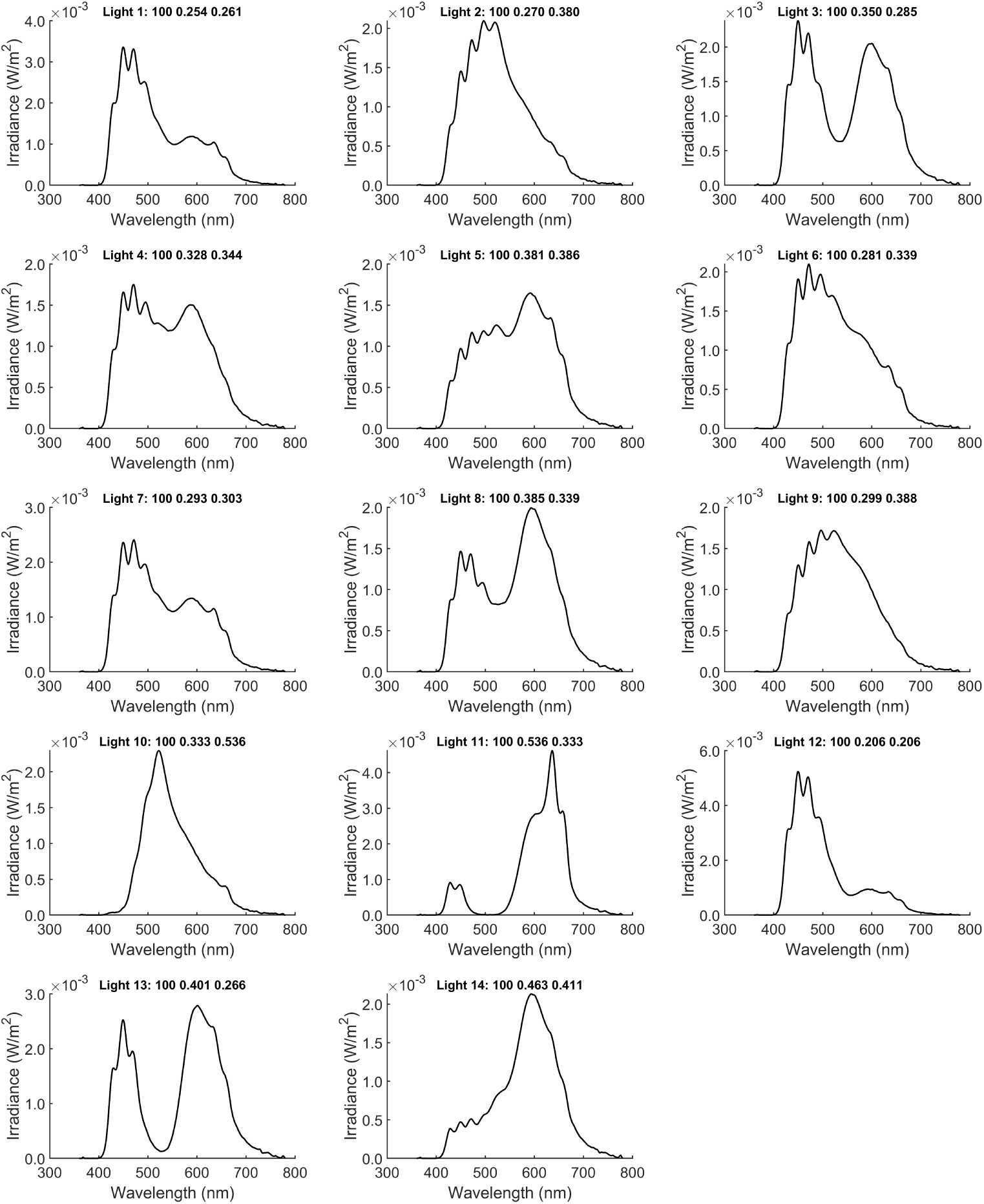
Illumination spectra. Spectra were generated as smooth metamers of the requested illumination chromaticities. Lights 10 through 14, of the “extreme” illumination subset, are relatively less broadband than the others, due to their higher saturation.

### 2.6. Surrounds

The observer’s visual field consisted of the adjustable display surface (the “patch”: 5° by 9° visual field) embedded within the black or grey immediate surround (54° by 55° and 44° by 45° for black and grey surrounds, respectively) and further enclosed within the backdrop of the lightroom white walls (filling the remainder of the visual field for both surrounds), as shown in 4a. The overhead lamps cast a gentle illumination gradient on both the walls and the cardboard surrounds. The lights being calibrated for irradiance at the observers’ eyes, the gradient affected only the local distribution of the total irradiance. Here we characterise the gradient on both the cardboard surround and the white walls. Both surrounds were of fairly flat spectral reflectance, as shown in Figure 3.

**Fig. 3.**
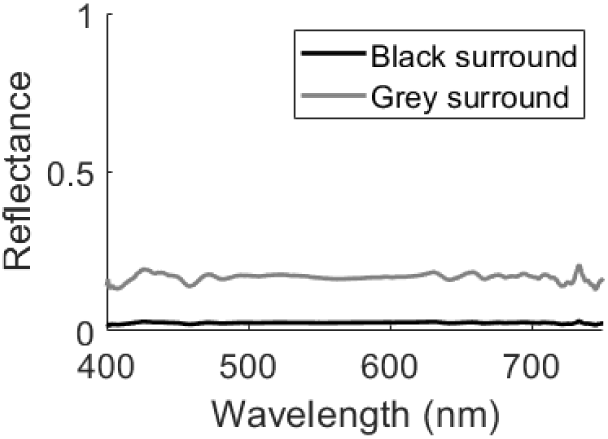
Surround reflectance spectral uniformity

Measurements of the surround boards (Table 2 and Table 3) were discretely sampled as a set of six points - top-left (TL), top-right (TR), immediate left of centre patch (CL), immediate right of centre patch (CR), bottom-left (BL) and bottom-right (BR). Measurements of the background white wall (Table 4) were sampled as a 3×3 uniform grid of evenly distributed measurement foci – top-left (TL), top-middle (T), top-right (TR), left (L), centre (C), right (R), bottom-left (BL), bottom (B), and bottom-right (BR). The luminance at the walls was higher in the middle and bottom than the top, the walls being at greater horizontal depth behind the overhang of the lights, while the luminance gradient on the cardboard surrounds was downward decreasing, the surround boards being at an acute vertical angle beneath the lights. The left and right fields of both the walls and the cardboard surrounds had negligible differences. The vertical luminance gradient on the backdrop of the walls ranged from a factor of 0.9 to 1.1 of the luminance at the centre of the wall (the centre of the wall being occluded by the surround-patch setup). The luminance gradient on the grey surround board flowed from a factor of 1.5 to 0.75, top to bottom, of the luminance near the centre of the board, with a maximum chromatic error between chromaticity measurements *m*_*i*_ and EEW of 3 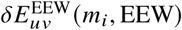 anywhere on its surface (and similar for 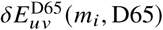 under D65 illumination). The low-reflectance black surround board shared a similar luminance gradient factor (1.5 to 0.75, top to bottom) with respect to its centre and dropped to a maximum of 2.15 cd/m^2^ anywhere on its surface. The chromatic error between chromaticity measurements *m*_*i*_ and EEW at any point on the black surround was within 3.5 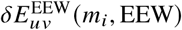 (and similar for 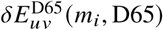 under D65 illumination). All surround measurements were done under illuminants which produced 100 lux of illuminance at the eye.

**Table 2.** Black surround spatial colorimetry

| Pos. | Y (cd/m<sup>2</sup>) | CIE x | CIE y | $\delta E_{uv}^{EEW}$ |
| --- | --- | --- | --- | --- |
| TL | 2.15 | 0.3357 | 0.3331 | 2.29 |
| TR | 2.06 | 0.3324 | 0.3291 | 3.37 |
| CL | 1.41 | 0.3374 | 0.3349 | 3.42 |
| CR | 1.26 | 0.3362 | 0.3342 | 2.44 |
| BL | 1.13 | 0.3370 | 0.3338 | 3.25 |
| BR | 1.21 | 0.3369 | 0.3341 | 3.09 |

**Table 3.** Grey surround spatial colorimetry

| Pos. | Y (cd/m<sup>2</sup>) | CIE x | CIE y | $\delta E_{uv}^{EEW}$ |
| --- | --- | --- | --- | --- |
| TL | 14.08 | 0.3307 | 0.3315 | 2.28 |
| TR | 13.10 | 0.3302 | 0.3303 | 2.97 |
| CL | 9.76 | 0.3310 | 0.3316 | 2.04 |
| CR | 9.78 | 0.3309 | 0.3315 | 2.14 |
| BL | 7.58 | 0.3320 | 0.3321 | 1.24 |
| BR | 7.81 | 0.3306 | 0.3309 | 2.51 |

**Table 4.** Background wall spatial colorimetry

| Pos. | Y (cd/m<sup>2</sup>) | CIE x | CIE y | $\delta E_{uv}^{EEW}$ |
| --- | --- | --- | --- | --- |
| TL | 27.21 | 0.3465 | 0.3436 | 11.45 |
| T | 27.03 | 0.3460 | 0.3431 | 10.99 |
| TR | 27.11 | 0.3465 | 0.3437 | 11.47 |
| L | 33.55 | 0.3427 | 0.3401 | 8.06 |
| C<sup>a</sup> | 30.44 | 0.3429 | 0.3397 | 8.13 |
| R | 33.47 | 0.3429 | 0.3403 | 8.24 |
| BL | 31.94 | 0.3350 | 0.3324 | 2.03 |
| B<sup>a</sup> | 31.58 | 0.3343 | 0.3320 | 1.73 |
| BR | 31.24 | 0.3351 | 0.3327 | 1.94 |

<sup>a</sup>occluded

We report 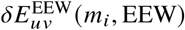 as the chromatic (*u,v* only) error, between the measured chromaticity *m*_*i*_ and the intended chromaticity (EEW), in the visual field. The chromatic error 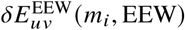 on the walls are larger, however the visual field consists mostly of the patch and surround board, the walls only being beyond 54° (black surround) and 44° (grey surround) in the peripheral visual field. The immediate surround of the central patch on the grey surround board closely matches the patch luminance of 10 cd / m^2^. The patch is 1.024 times brighter (0.024 Weber contrast) than the grey surround, and 7.491 times brighter (6.491 Weber contrast) than the black surround.

### 2.7. Experiment procedure

Prior to each experiment session, the lamps were switched on at EEW with experiment luminance for thirty minutes, and the tablet was inserted into its slot in the wooden frame. The surround board (grey or black) for the session was attached to the Velcro strips on the frame. At experiment start, the participant was draped with a black cloth over their shoulders and seated at the chinrest, any necessary chinrest and frame tray height adjustments then being carried out. A black and white printed sheet with the four navigation colour directions (up for yellow, down for blue, left for green, right for red) was placed on the floor, out of the field of view from the observing position at the chinrest, but to which participants could refer by briefly deviating their head from the chinrest. Instructions for the adjustment task were verbally provided for each participant’s first session and reconfirmed for subsequent sessions, including confirmation of participant understanding of the colour navigation directions, and the controller buttons for navigation and indicating satisfactory matches. Between trials, participants were asked to look at some arbitrary spot on the surround board, during which time they could stretch their neck, if required, by briefly removing their head from the chinrest. Participants were allowed an optional break of 5 minutes halfway through each session of fourteen illuminations. If they chose to take the break, the control program was paused, the participant brought out of the lightroom into the low illumination exterior chamber, reseated at the chinrest at the end of the break, and the control program resumed. To reset the participant’s adaptation state, each new block of trials began with 120s exposure to the baseline (reference) white light, D65.

The experiment control program (Appendix A) was then launched. Each experiment session tested of fourteen ambient illuminations for one surround, with six achromatic adjustment trials per illumination. Each set of six trials lasted a total of 460s. For each illumination, a leading adaptation period of 120s under D65 was followed by an alternating series of 30s adaptation to the illumination and 25s of device screen adjustment (35s for first trial). The starting chromaticity for the first trial in the set of six was set to a random point _*i*_ within 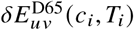 from the test illumination chromaticity *T*_*i*_, and for each subsequent trial was set to the endpoint of the previous trial. During the 30s adaptation-only periods, the device screen chromaticity was reset to match the ambient illumination. Throughout the set of trials, the ambient illumination remained static, providing a continuously increasing duration of exposure. See Figure 4 for the setup diagram and timeline of trials.

**Fig. 4.**
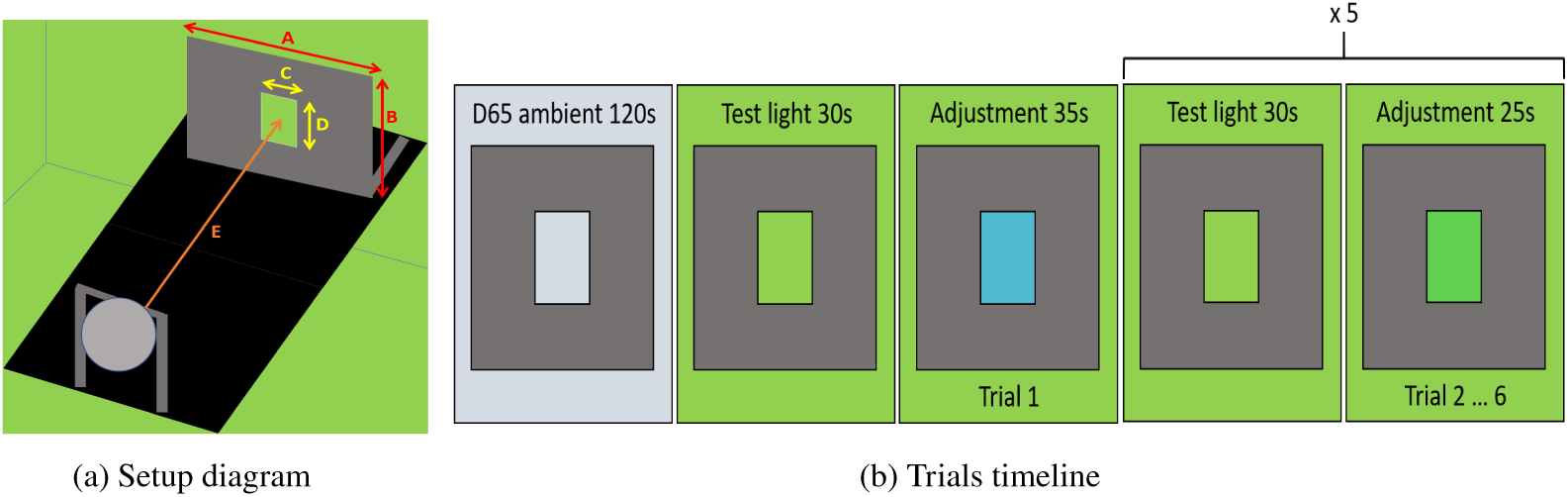
(a) Setup diagram and (b) trials timeline. Setup diagram: A, B surround dimensions. C, D patch dimensions. E viewing distance. Trials timeline: Sequence of trials per illumination. Trial 1: Patch set to random chromaticity. Trials 2 to 6, adjustment proceeds from endpoint of previous trial. 30 seconds of top-up adaptation between trials, with the display chromaticity set to match the ambient illumination chromaticity at the start, and reset to the endpoint of the previous trial at the end.

Participants were asked to mark any chromaticities that they perceived as satisfactorily neutral, at any point within each 25s adjustment slot (35s for first slot), by pressing an auxiliary controller button, to which the control program would respond with the auditory signal “happy”. Participants could mark satisfactory matches multiple times during a trial, and moreover to continue making screen chromaticity adjustments even after indicating a satisfactory neutral, as long as it was within the time bounds of the adjustment slot. Between adjustment slots, the controller button for indicating satisfactory matches and the navigation buttons were disabled. We hereafter refer to chromaticities marked satisfactory neutral as “confirmed matches”. Input times for all confirmed matches were recorded.

### 2.8. Ethics

The study was reviewed and approved by the Faculty of Medical Sciences Research Ethics Committee, part of Newcastle University’s Research Ethics Committee (approval 1520/6052), which includes members internal to the Faculty, as well as one external member. Committee members must provide impartial advice and avoid significant conflicts of interests. Written consent was received from all participants prior to participation in the study.

## 3. Analysis and results

### 3.1. Analysis

The data consists of chromaticity adjustment trajectories over six trials per participant per illumination. The adjustment end-point in each trial is the “navigational” match, while any adjustment trajectory chromaticities marked as satisfactory are the “confirmed” matches, with the latest confirmed match in each trial being the “trial-final” match. For both navigational and confirmed matches, the trial-final match on the sixth trial is the “illumination-final” match. For both navigational and confirmed matches, the data was analysed in terms of: (a) surround – trial-final matches over all trials, illuminations and participants, for each surround; illumination-final matches over all illuminations and participants, for each surround, (b) illumination – trial-final matches over all trials and participants, for each illumination and each surround; illumination-final matches over all participants, for each illumination and each surround, (c) time – trial-final matches over all illuminations and participants, for each trial (both trial number and adaptation offset) and each surround.

A colour constancy index (CCI) was calculated, for individual and mean matches, as an indication of completeness of chromatic adaptation. The CCI (Equation 1) takes the ratio of the distance between the achromatic point and the test illumination to the distance between the reference (baseline) illumination and the test illumination chromaticities, in CIELUV colour space, and subtracts this from 1. Partial adaptation towards the test illumination is indicated by values within the range 0 to 1 (negative values indicate matches shifted away from the adapting illumination instead of towards it). An extension to the CCI, the projected CCI, or pCCI (Equation 2), takes the projection (in CIELUV colour space) of the vector from the achromatic point to the test illumination on the line from the test illumination to the reference illumination. It discounts lateral adjustments and expresses constancy only along the axis connecting the reference and test illumination chromaticities (Figure 5).

**Fig. 5.**
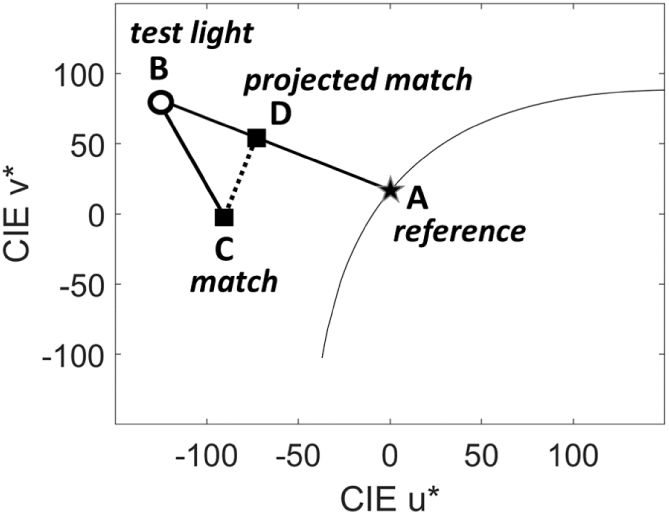
Colour constancy (1 − *BC* /*AB*) and projected colour constancy (1 − *BD* /*AB*) indices in CIELUV colour space. To calculate the projected colour constancy index *pCCI*, the achromatic match is projected onto the line between the test illumination and the reference illumination chromaticities. The black curve represents the daylight locus.

Let *A* be the reference light chromaticity, *B* the test light chromaticity and *C* the match chromaticity, in CIELUV colour space. The CCI is given by

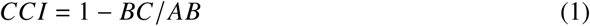

where *BC* is the distance between the achromatic point and the test illumination chromaticities, and *AB* the distance between the reference illumination and the test illumination chromaticities.

We define the “projected match” *D* as the match coordinates projected onto the line joining the test light and the reference light chromaticities in CIELUV colour space. The pCCI is then given by

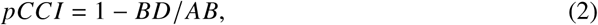

where *BD* is the distance between the projected match and the test illumination chromaticities.

The 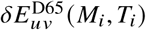 of the match or projected match is an absolute measure of discrepancy between the match *M*_*i*_ and test chromaticity *T*_*i*_. The CCI and pCCI are the corresponding relative measures of match goodness.

The statistical analyses of achromatic match data wese conducted using MATLAB (version R2018a, MathWorks, Natick, MA, USA). N-way repeated measures analyses of variance (ANOVA) were run with fixed factors of light condition (14 lights), surround type (black, grey), and timepoint (initial, final); and participants as a random factor (MATLAB function anovan). Multiple comparisons of estimated marginal means between factors and factor levels were also performed (MATLAB function multcompare, invoked with Bonferroni adjustment). All factors were coded as categorical variables, with only the dependent variable being left in its native data format (floating point for CCI and pCCI data). In separate analyses, parameters of the adaptation progression model (described in the following section), as well as quantifiers of illumination properties, were compared using paired-sample T-tests (MATLAB function ttest), as well as Pearson’s correlations (MATLAB function corrcoef) for which *p* values were Bonferroni-adjusted for multiple comparisons.

Note that in statistical analyses and plots, only the final navigational or confirmed match at each timepoint was used, even though participants had been allowed to make multiple match submissions (in case they felt they could do better, after performing navigation or submitting a match, within any remaining time during a time slot). Final navigational and confirmed matches superceded earlier ones at each timepoint.

### 3.2. Results

Achromatic adjustment trajectories typically consisted of varying numbers of controller inputs per trial, with decreasing number of inputs from trial 1 to trial 6. Early trials have a larger traversal of CCI, and pCCI, and require more controller inputs. Later trials have a smaller change in CCI, and pCCI, and require fewer controller inputs, indicating that the distances *BC* and *BD* (Equations 1, 2 and Figure 5) change by decreasing amounts as trials progress. Changes in CCI and pCCI were thus proportional to the number of controller inputs, as observed by the linearity of CCI and pCCI progression by navigational input number in Figure 6, which shows examples of adjustment trajectories and levels of adaptation, as indicated by the CCI and pCCI metrics, at each navigation step. The proportionally decreasing magnitude of CCI and pCCI changes with time is symptomatic of a proportional rate progression of adaptation, which we fit with an exponential model. Later trials also typically had a larger number of submitted matches, also due to proportional rate progression as smaller changes are required to achieve satisfactory matches as adaptation, mirrored by adjustment, proceeds.

**Fig. 6.**
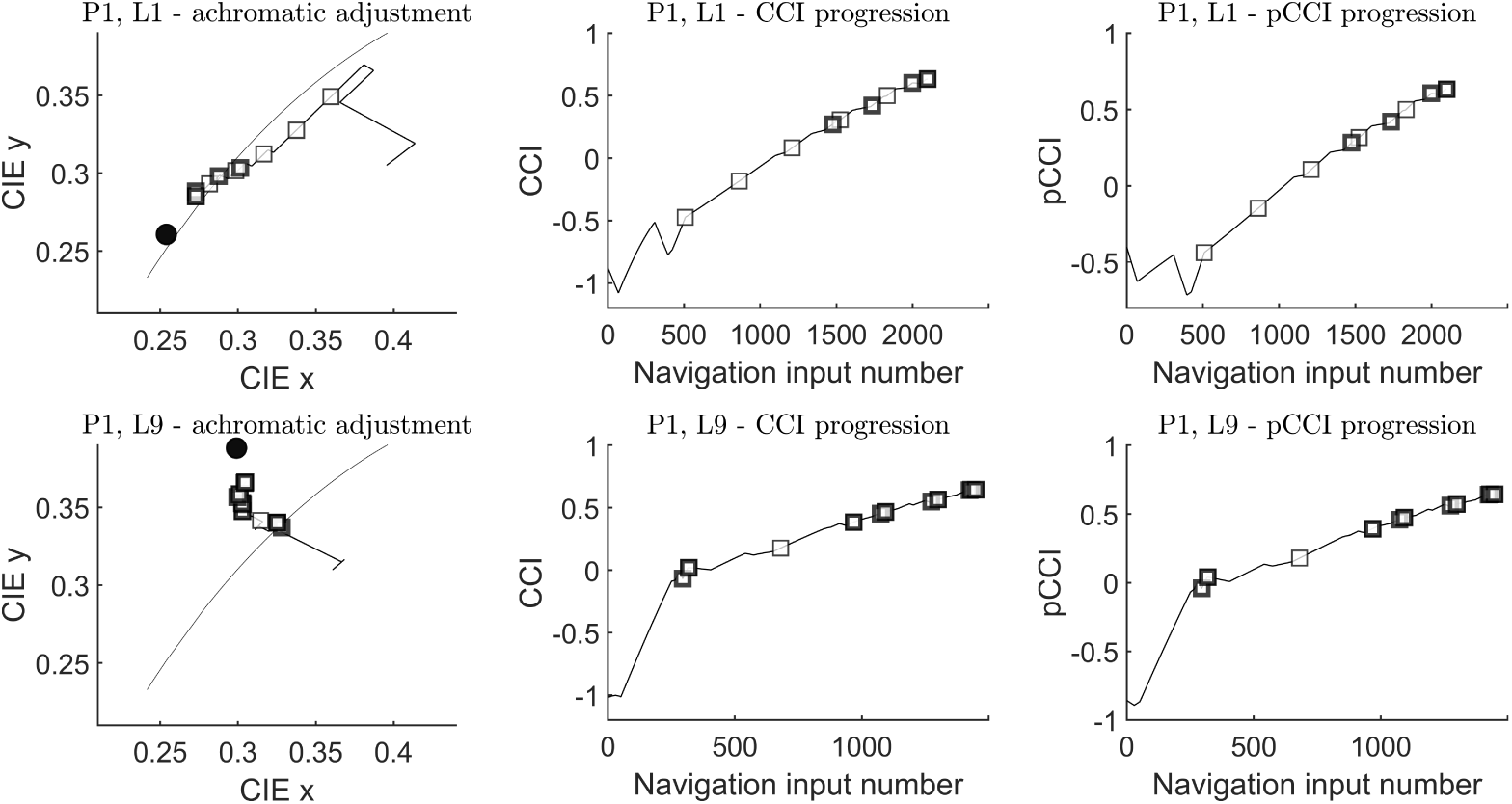
Examples of achromatic adjustment trajectories, CCI progression, and pCCI progression (black surround), by navigation step. Black circles (left plots) indicate test illumination chromaticity. Light and dark squares indicate trial-final navigational matches and trial-final confirmed matches, respectively. The navigation input number axis shows the varying number of discrete navigation inputs via the controller, per instance of achromatic adjustment, within each adjustment slot as well as over all six trials.

Adaptation, indicated by the colour constancy indices CCI and pCCI, improved over time, from initial measurements between 30 and 65 seconds to endpoints between 315 and 340 seconds, for all lights and both surrounds. Colour constancy indices for the grey surround were higher than for the black surround (mean matches per light for each surround illustrated by Figure 7), for all lights and at both initial and final timepoints (Figure 8). Although the indices differed between lights, there was no systematic difference between lights on and off the daylight locus, nor between lights of moderate and extreme chromaticity. The time course of adaptation, as measured by migration of the pCCI, differed between lights and between participants. Adaptation appears to continue beyond 340 seconds (5 minutes and 40 seconds).

**Fig. 7.**
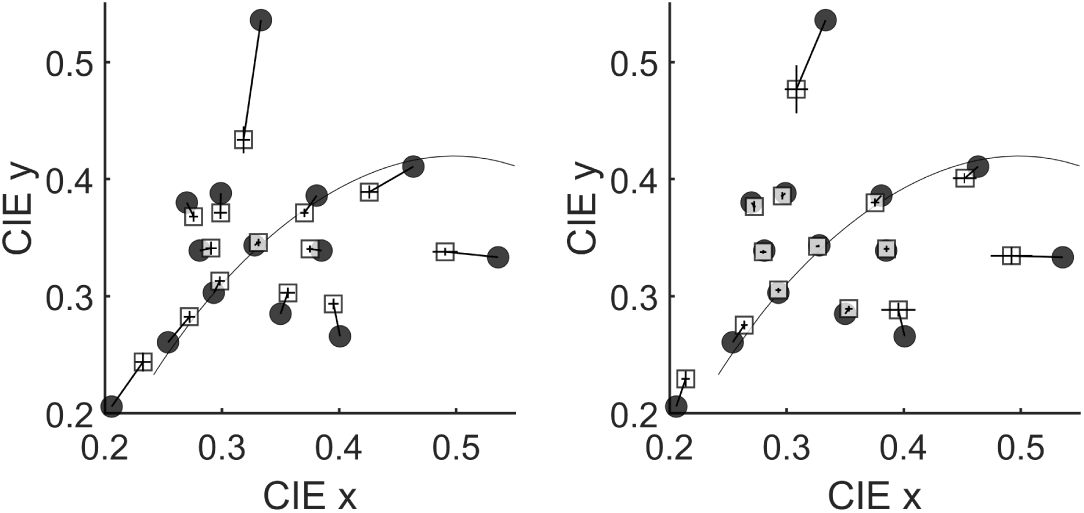
Left: black surround mean achromatic matches. Right: grey surround mean achromatic matches. Circles indicate test illumination chromaticities. Squares indicate mean matches over all participants, with error bars as the standard errors of mean CIE x and mean CIE y coordinates.

**Fig. 8.**
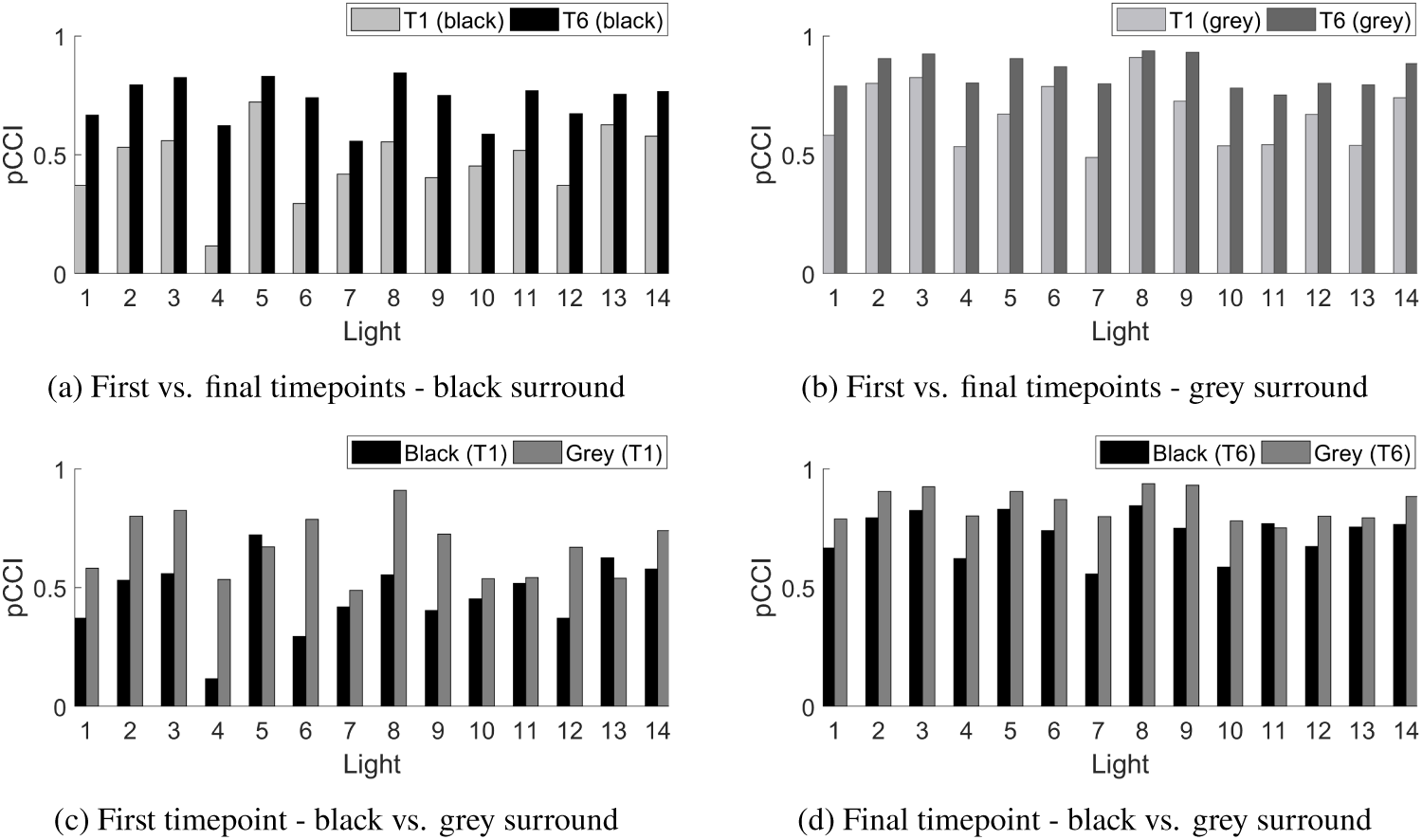
Mean pCCI values over all participants between (a) first and final timepoints - black surround, (b) first and final timepoints - grey surround, (c) first timepoint for black surround and first timepoint for grey surround, (d) final timepoint for black surround and final timepoint for grey surround.

Repeated measures ANOVAs, with fixed factors of light, surround and timepoint, confirm significant interactive and main effects of the three factors (see Table 5 for F-statistics and *p* values). Comparisons of estimated means reveals that navigational CCIs and pCCIs for the grey surround are significantly higher than those for the black surround (mean difference 0.128, *p <* .001 and 0.126, *p <* .001, respectively), and significantly higher for the final timepoint than for the first (mean difference 0.363, *p <* .001 and 0.313, *p <* .001, respectively).

**Table 5.** Main (light, surround, timepoint) and interactive (light*timepoint, surround*timepoint) effects from repeated measures ANOVA over navigational match CCI (N. CCI) and pCCI (N. pCCI), and confirmed match CCI (C. CCI) and pCCI (C. pCCI) values. Only significant effects reported.

|  | Light | Surround | Timepoint | Light*Timepoint | Surround*Timepoint |
| --- | --- | --- | --- | --- | --- |
| N. CCI | $F(13, 91.24) = 7.69$<br>$p < .001$ | $F(1, 7.00) = 11.26$<br>$p = .01$ | $F(1, 7.00) = 145.5$<br>$p < .001$ | $F(13, 291.0) = 2.32$<br>$p = .01$ | - |
| N. pCCI | $F(13, 91.23) = 5.66$<br>$p < .001$ | $F(1, 7.00) = 13.60$<br>$p = .01$ | $F(1, 7.00) = 93.21$<br>$p < .001$ | - | - |
| C. CCI | $F(12, 112.8) = 7.37$<br>$p < .001$ | $F(1, 7.87) = 26.23$<br>$p < .001$ | $F(1, 7.47) = 42.56$<br>$p < .001$ | $F(13, 197.0) = 2.54$<br>$p = .003$ | $F(1, 197.0) = 5.83$<br>$p = .02$ |
| C. pCCI | $F(12, 113.3) = 4.52$<br>$p < .001$ | $F(1, 8.33) = 31.10$<br>$p < .001$ | $F(1, 7.44) = 25.21$<br>$p = .001$ | - | $F(1, 197.0) = 5.61$<br>$p = .02$ |

We model the temporal progression of adaptation by a proportional rate growth function [33,34], given by

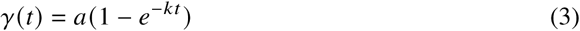

where *γ* (*t*) is adaptation level (in our case, CCI or pCCI) at time *t* (seconds), and *a* and *k* are free parameters.

The speed of adaptation progression is regulated by the factor *k*. In general, progression consists of faster growth for higher values of *k*, approaches linear growth as *k* approaches 0, and switches to decay for negative *k*. Parameter *a* determines the asymptote of progression and therefore describes the upper limit, or extent, of adaptation (for each data subset), with adaptation level *γ* approaching *a* for large *t*.

We evaluate goodness of fit using Matlab’s ‘goodnessOfFit’ function, with the normalized root mean square error (NRMSE) cost function. Its values range from negative values (poor fit), through 0 (no better than a straight line fit), to 1 (perfect fit). Where fitting the model to pooled data to obtain summary model parameters by condition or participant subsets, the goodness of fit is evaluated against timepoint (adjustment slot) means. For navigational matches, this is the end of the slot, while for submitted matches, the timepoint mean is the mean time of submission of all the submitted matches recorded within the slot. Pooled data has higher scatter due to data from multiple conditions or participants (with high inter-condition and inter-participant differences) being combined. As we go on to discuss, the proportional rate growth model nonetheless represents well the adaptation values at timepoint means over various subsets of pooled data, with differences in the free parameters *a* and *k* characterising differences in adaptation between the subsets. Model fits to pCCI data for all lights and participants are given in Appendix B.

Globally fitting the growth function to all confirmed match pCCI data pooled together for the black surround (Figure 9a) and then separately for the grey surround Figure 9b, we find *a* = 0.725, *k* = 0.016 (fit 0.934) for the former, and *a* = 0.867, *k* = 0.026 (fit 0.938) for the latter. Taking the mean^5^ of parameter values over all lights and then for all participants per surround also yields similar values - mean over lights for black surround: *a* = 0.726, *k* = 0.019 (mean fit 0.835); mean over participants for black surround: *a* = 0.740, *k* = 0.024 (mean fit 0.868); mean over lights for grey surround: *a* = 0.873, *k* = 0.028 (mean fit 0.889); mean over participants for grey surround: *a* = 0.855, *k* = 0.029 (mean fit 0.884). Parameter means over all lights are a better predictor of the adaptation progression growth function parameter values for the pooled data, than are parameter means taken over all participants, due to inter-participant variability. For the grey surround, parameter means over both lights and participants better match the pooled fit, indicating lower inter-participant variability, than they do for the black surround. Model parameters for all fourteen lights are given in Table 6, and those for all participants in Table 7.

**Fig. 9.**
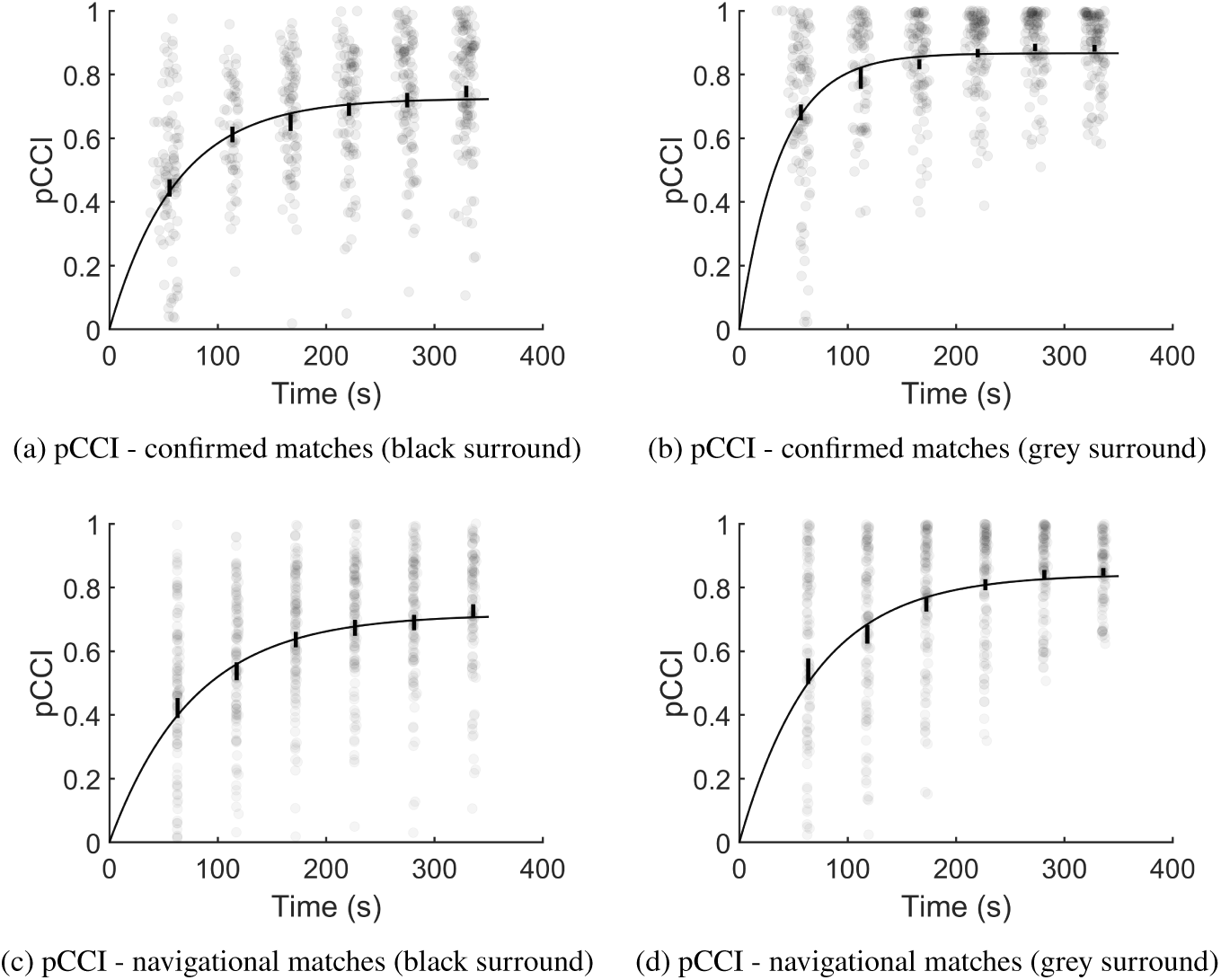
Adaptation progression growth function fits to pCCI data pooled over all lights and participants, for black and grey surround conditions. Confirmed matches are adjustment chromaticity coordinates marked as satisfactorily neutral by the participant. Navigational matches are adjustment chromaticities from trial end-points. Grey markers indicate individual matches at each of trials 1 through 6. Error bars: standard errors of timepoint means.

**Table 6.**
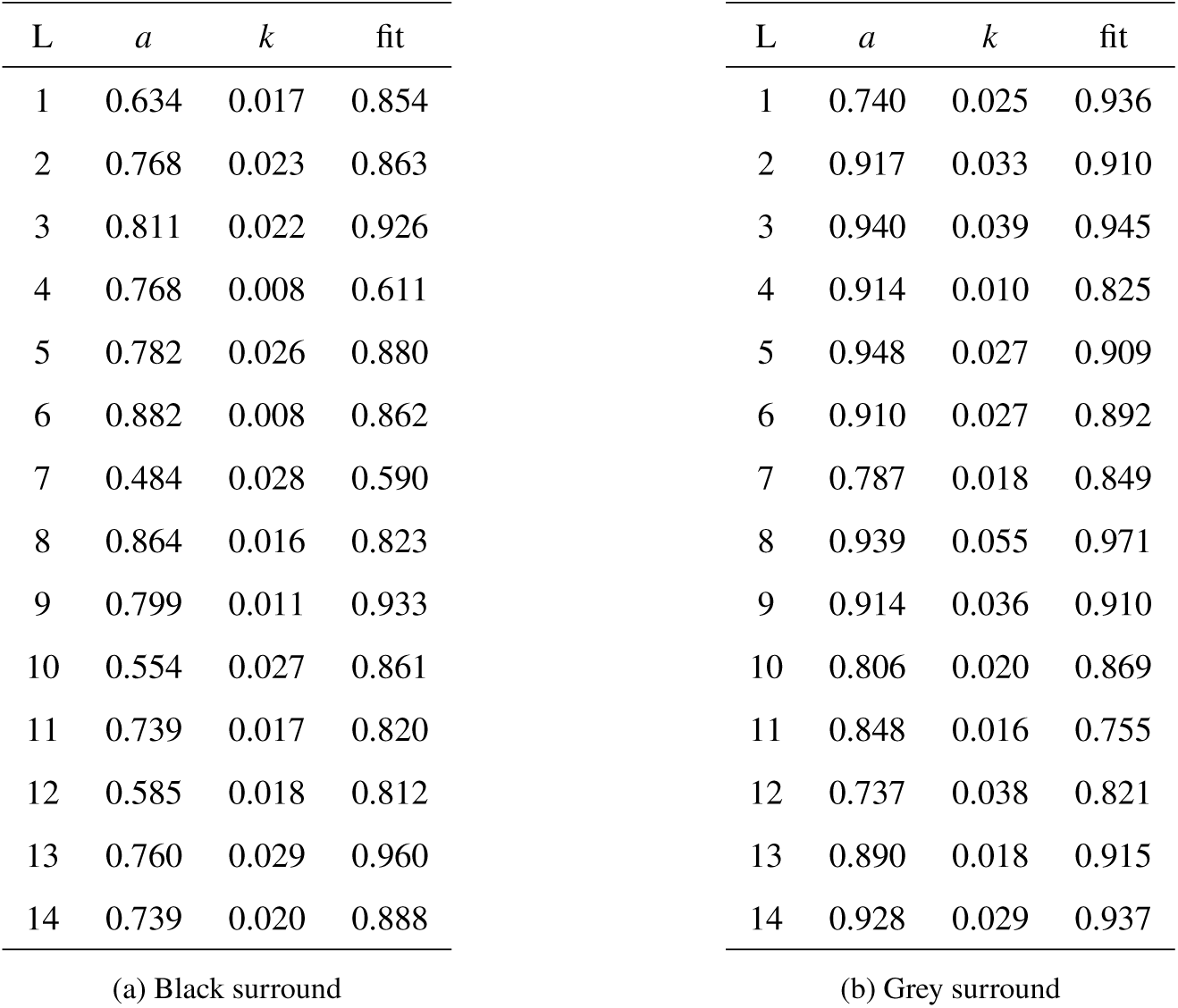
Proportional rate growth parameters *a* and *k* for adaptation progression, and goodness of fit, per light (confirmed match pCCIs pooled across participants). Daylight locus lights: mean *a* = 0.681, *k* = 0.020 (mean fit 0.764) for black surround, and *a* = 0.864, *k* = 0.022 (mean fit 0.891) for grey surround. Off-locus lights: mean *a* = 0.751, *k* = 0.019 (mean fit 0.873) for black surround, and *a* = 0.878, *k* = 0.031 (mean fit 0.888) for grey surround. Over all lights, mean *a* = 0.726, *k* = 0.019 (mean fit 0.835) for black surround, and *a* = 0.873, *k* = 0.028 (mean fit 0.889) for grey surround.

**Table 7.**
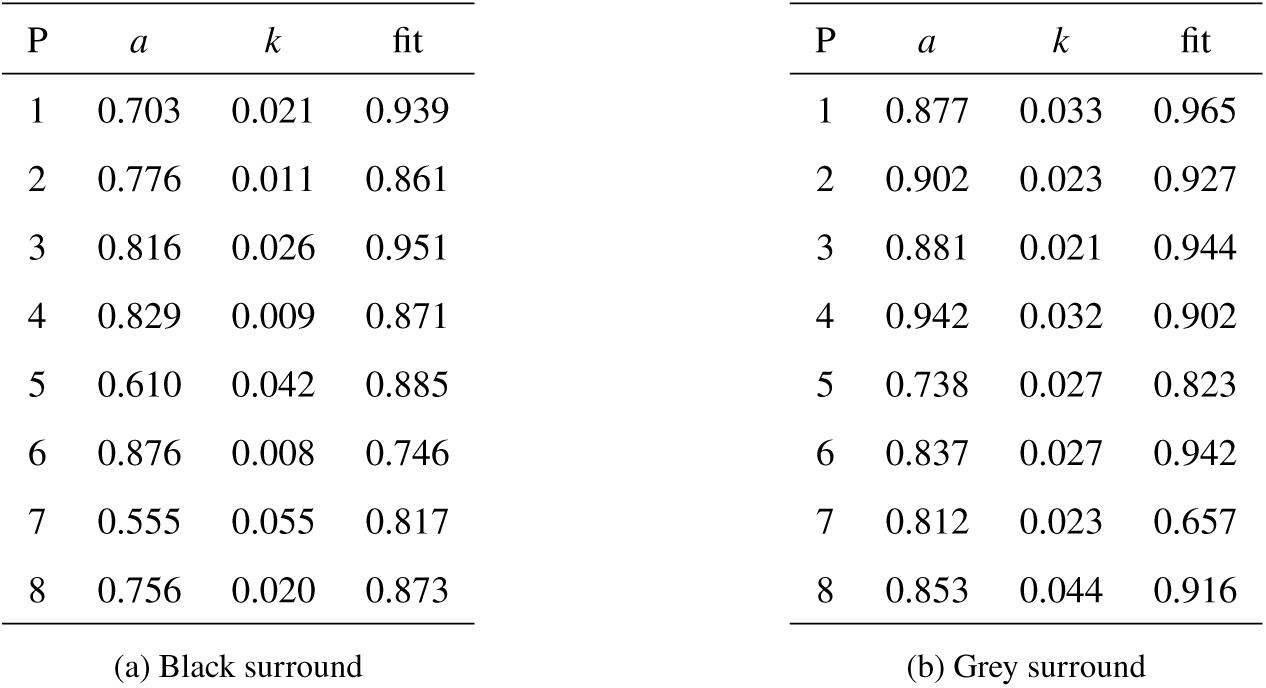
Proportional rate growth parameters *a* and *k* for adaptation progression, and goodness of fit, per participant (confirmed match pCCIs pooled across lights). Participant mean *a* = 0.740, *k* = 0.024 (mean fit 0.868) for black surround, and *a* = 0.855, *k* = 0.029 (mean fit 0.884) for grey surround.

Over all lights, in paired-sample T-tests, parameter *a* (*t* (13) = −7.98, *p <* .001), parameter *k* (*t* (13) = −2.27, *p* = .041), and goodness of fit (*t* (13) = −2.17, *p* = .049), are significantly larger for the grey surround versus the black surround (Figure 10). The larger *a* and *k* values for the grey surround indicate better extent and rate of adaptation over the test lights. Over all participants, only parameter *a* (*t* (7) = −3.83, *p* = .006) is significantly larger for the grey surround, although the data for *k* and goodness of fit also trend towards being higher.

**Fig. 10.**
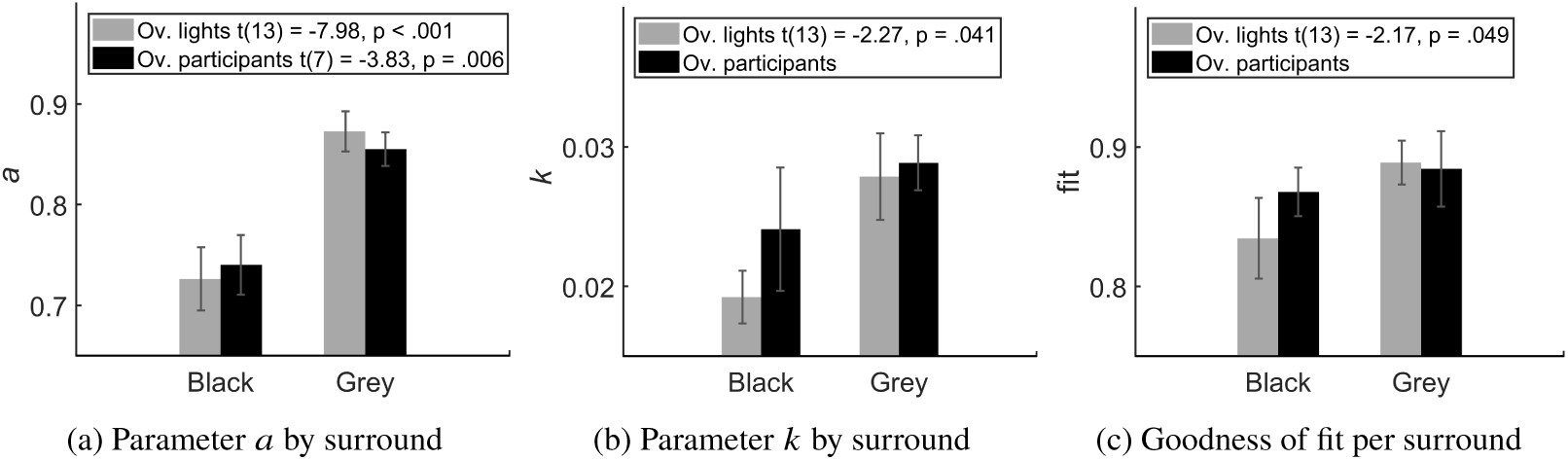
Mean model parameters *a* and *k*, and goodness of fit, by surround (from confirmed match pCCI data), calculated over lights and over participants. Error bars: standard errors of means. Parameter *a* is significantly higher for the grey surround, both over lights and participants, indicating better extent of adaptation. Parameter *k* is significantly higher only over lights, as is goodness of fit.

Adaptation extents *a*, over all test lights, are strongly correlated between black and grey surrounds (*r* (12) = .84, *p <* .01), as are 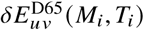 of mean confirmed match *M*_*i*_ from test light *T*_*i*_ (*r* (12) = .92, *p <* .001). Adaptation rates *k* over the test lights are not significantly correlated between the two surrounds, even though its values are significantly different between the surround types as identified by the previously mentioned T-tests (Figure 10). Interestingly, despite the lack of significant correlation between surrounds for fit parameter *k*, its value for light 4 is very low for both surrounds (Table 6a and Table 6b) compared to its value for other lights, suggesting that adaptation to light 4 is relatively slow, which may be explained by light 4 being nearest both EEW and the reference D65, and the evidence for a range of neutral chromaticities being accepted as “white” by observers [35,36].

Overall, extreme illuminations appear to produce similar adaptation progression rates and extents as moderate illuminations. The same holds for illumination chromaticities on and off the blackbody and daylight loci. Undifferentiated adaptation extent for different chromaticities of reference illumination translates to larger absolute errors in matches for extreme chromaticities, confirmed by the strong correlation between extremeness 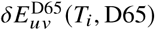 of test light chromaticity *T*_*i*_ with respect to the baseline illumination (D65) and the distance 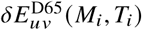 of mean confirmed match *M*_*i*_ from the test light *T*_*i*_, for both black (*r* (12) = .91, *p <* .001) and grey (*r* (12) = .95, *p <* .001) surrounds (Figure 11). We find no general dependence of adaptation extent and rate on illumination saturation, with adaptation extent *a* and pCCI for both surrounds correlating weakly with illumination extremeness as judged with respect to the reference D65. Interestingly, for the black surround, 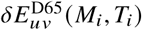 is better correlated with illumination extremeness 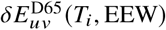 with respect to EEW (*r* (12) = .94, *p <* .001) than with extremeness 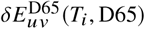 with respect to the baseline reference D65 itself (*r* (12) = .91, *p <* .001), but are similar for the grey surround.

**Fig. 11.**
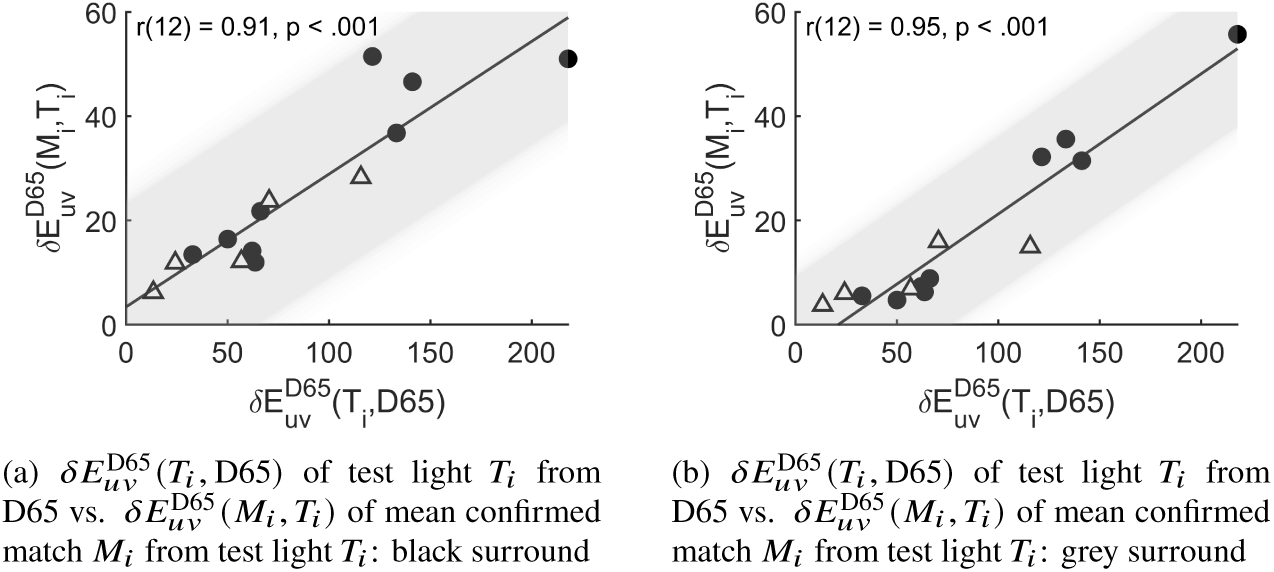
Correlations (Pearson’s *r*) between chromatic difference 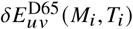 of mean confirmed match *M*_*i*_ from test light *T*_*i*_ against chromatic difference 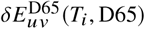 of test light *T*_*i*_ from D65. Triangles indicate lights on the daylight locus.

The pCCI progression curve, and therefore the proportional rate growth model fit, is variable between participants and between lights (Appendix B). This of course means that the overall mean model parameters are of limited specific predictive power, without further extension considering other factors possibly including characteristics of the ambient light, immediate surround and individual differences. However, the goodness of fit with two degrees of freedom for individual progression curves, over different lights and participants, is a confirmation of the proportional rate growth model as a representation of adaptation progression.

## 4. Discussion

We measured the progression of adaptation under 14 distinct real illuminations, in a naturalistic immersive viewing environment, tracking the change in participants’ perceptual whitepoints – measured by achromatic adjustment of a small test patch - over time as they adapted to each distinct illumination condition. We held the illumination reaching the eye constant while varying the local surround of the adjustable test patch, across two different conditions, one a neutral reference surface, and the other black. Thus we were able to disentangle spatial contrast effects from temporal adaptation to the ambient illumination, and thereby address questions arising from differences in previously reported studies.

Adaptation level was quantified by the relative closeness of the achromatic setting to the adapting illumination chromaticity, using colour constancy indices, with a range varying from negative values to 1, where 1 marks an exact match between the two. Colour constancy indices (CCIs) improved over time, up to the maximum tested duration of 340 seconds, for all lights and both surrounds. CCIs were higher for the grey surround than for the black, for all lights and at all timepoints. Although the timecourse of adaptation varied between lights, its endpoint varied little, with the projected CCI (pCCI) reaching a mean of 0.85 for the grey and 0.73 for black, across all lights.

We modelled the progression of chromatic adaptation, and the consequent change in colour constancy, using a proportional rate growth function. Growth functions cannot straightforwardly be characterised by a static-valued half-life, so we instead refer to the asymptote limit *a* and the growth factor *k*. We find that *a* and *k* vary between lights, with typical ranges of 0.5 to 0.95 and 0.01 to 0.06, respectively. The inter-participant variability in these parameters is larger. The time required to reach a pCCI of 0.5 – a pseudo-half-life - is about 60 seconds for the black surround and 25 seconds for the grey surround. The latter value agrees with previous findings [16, 17], in particular with the reported half-life of approximately 20 seconds for the “slow” phase of short-term adaptation [17].

The slower rate and lower extent of chromatic adaptation for the black versus the grey surround (both *a* and *k* are smaller for the former) supports previous findings of the effects of changes in the test stimulus or background intensity [22, 24, 26], and allows us to conclude more generally that decreasing contrast between the test stimulus and its background improves accuracy of the achromatic settings, i.e. increases the closeness of the perceptual whitepoint to the adapting light chromaticity. The fact that the difference between surrounds is pronounced even at the first timepoint, after 30 seconds of adaptation, suggests that the improvement in constancy is due to very fast processes, consistent with the instantaneous spatial contrast phase of adaptation described previously [17,18]. The fact that we maintained constant illuminance and chrominance at the eye across the two surround conditions means that the difference in constancy is not due to global adaptation processes.

The high level of adaptation attained at the final timepoint with the grey surround is also consistent with the level reported by [8] for a similar immersive viewing setup, but with uncontrolled adaptation durations. One question is whether the presence of recognizably neutral surfaces in the variegated scene [8] served as reference surfaces to which participants might make direct matches. In the current study, the same may be asked of the grey surround. But if this were the case, that grey surfaces within the scene serve as direct references, then all achromatic settings, even those early on in the series of six trials, would correspond closely to the illumination colour, and there would be no gradual progression of the match towards the test illumination chromaticity. Yet, for both surround conditions, there is a significant improvement in constancy over time, and although the presence of a comparable luminance reference surface does not collapse the adaptation process to a short-lived event, it does facilitate its progression.

The main factor affecting deviation of the achromatic setting, we find, is the distance of the adapting illumination chromaticity from the baseline neutral illumination light (D65), to which all participants pre-adapted before exposure to each test illumination. The further the adapting illumination chromaticity from the neutral reference, the greater the absolute deviation of the achromatic setting. Correspondingly, the growth factor *k* does not vary systematically with the distance of the illumination chromaticity from neutral. Thus, extreme illuminations require a longer exposure duration to reach adaptation stability, perhaps expected from the larger difference between neural gains under larger test-reference illumination chromaticity differences. For certain illuminations and observers, adaptation takes longer than 5 minutes and 40 seconds to asymptote. The CCI (and pCCI) measure the deviation relative to the test-reference distance; therefore the asymptotic values for these indices vary little between test illuminations. In particular, we find no significant difference in constancy between illuminations on and off the daylight locus, nor for illuminations with moderate versus extreme chromaticities relative to the neutral reference chromaticity.

The proportional growth model is a useful, concise quantitative description of the progression of chromatic adaptation and colour constancy under steady global illuminations over a range of chromaticities. The goodness of fit for the model is generally high, with only two degrees of freedom (parameters *a* and *k*). These parameters do, nonetheless, vary between participants and between lights (Appendix B), implying that their mean values cannot be used universally to predict adaptation progression for arbitrary illuminations, viewing conditions, or individuals. Further exploration of the relationship between those factors and the growth function is needed. The correlation of the adaptation extent *a* between surrounds suggests that adaptation completeness is largely determined by illumination chromaticity, albeit not necessarily by illumination saturation. There is no corresponding correlation between the two surrounds for adaptation rate *k*, but we note that *k* has more variance over all test lights, with extreme illumination chromaticities also producing a less accurate model fit, for the grey surround. These differences may also explain the observation that although higher adaptation extents appear to indicate slower adaptation rates for the black surround, they do not for the grey surround.

## 5. Conclusions

In summary, we find that for immersive exposure to a chromatic illumination following pre-adaptation to neutral, the change in adaptation state over time (quantified by the relative closeness of the perceptual whitepoint to the test illumination chromaticity) (a) can be modelled by a proportional rate growth function, typically requiring more than 5 minutes to stabilise, (b) depends on the contrast between the test surface and its background, specifically increasing with decreasing test-background contrast and (c) is generally similar in both extent and rate for different test illumination chromaticities. Adaptation progression does not differ significantly between illuminations on or off the daylight locus, or between moderate versus extreme test illumination chromaticities. This study finds no significant chromatic biases in magnitude and direction of chromatic adaptation within the experimental paradigm used.

While chromatic adaptation is not synonymous with colour constancy, it is one of the primary underlying mechanisms, and likely the most powerful mechanism at the sensory level. Achromatic settings - the chromaticity perceived as neutral - provide a measure of adaptation to the ambient illumination, and in turn, a measure of colour constancy. Our results highlight important considerations when using achromatic settings as measures of colour constancy, especially when comparing between studies or illuminations. First, the reference illumination chromaticity must be taken into account, as its distance from the test illumination chromaticity will influence the extent of adaptation. Second, exposure duration to test illuminations must be systematically controlled, as adaptation progresses over time, in some cases beyond 5 minutes. Differences in colour constancy may therefore be partly due to differences in stimulus configurations, reference illumination chromaticities, and timepoints of measurements.

## 6. Acknowledgments

We thank the following for financial support: European Commission (H2020-MSCA-ITN DyViTo - grant 765121) and HI-LED – grant 619912; Huawei Device Co., Ltd. (R&D Agreement).

## Appendix A Experiment control program

The control program executed the following steps:

1. White adaptation. Set the ambient illumination to D65 for 120s. Set the LEDs to a new randomly chosen illumination from the set of test illuminations. Set trial counter to 1. Disable controller inputs.
2. Adaptation to test light for 30 seconds.
3. Adjustment trial.
  a. Set the device screen to a random chromaticity *c*_*i*_ within 300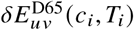 from the test light chromaticity *T*_*i*_ for the first trial, or to the last adjustment chromaticity for all other trials in each block.
  b. Issue the verbal instruction “first trial” only for trial 1, and “begin adjustment” for all trials, via speakers. Enable controller inputs.
  c. Monitor controller button-presses and record resulting chromaticity coordinate changes and relevant time offsets from start of adaptation to test light, for 35s at trial 1 or for 25s for trials 2 to 6.
  d. Issue the verbal instruction “stop” and reset the device screen to illumination chromaticity. Disable controller inputs. Increment trial counter.
  e. If trial counter is at 6, go to next step, else repeat steps 2 and 3.
4. Repeat steps 1 through 3, for each of the 14 illuminations, then issued the verbal notification “end of experiment”

The software interface allowed the experimenter to initiate experiment pause at any time during the trials, upon which the control program would pause before, and on request resume from, step 1.

## Appendix B pCCI progression for all lights and participants

**Fig. 12.**
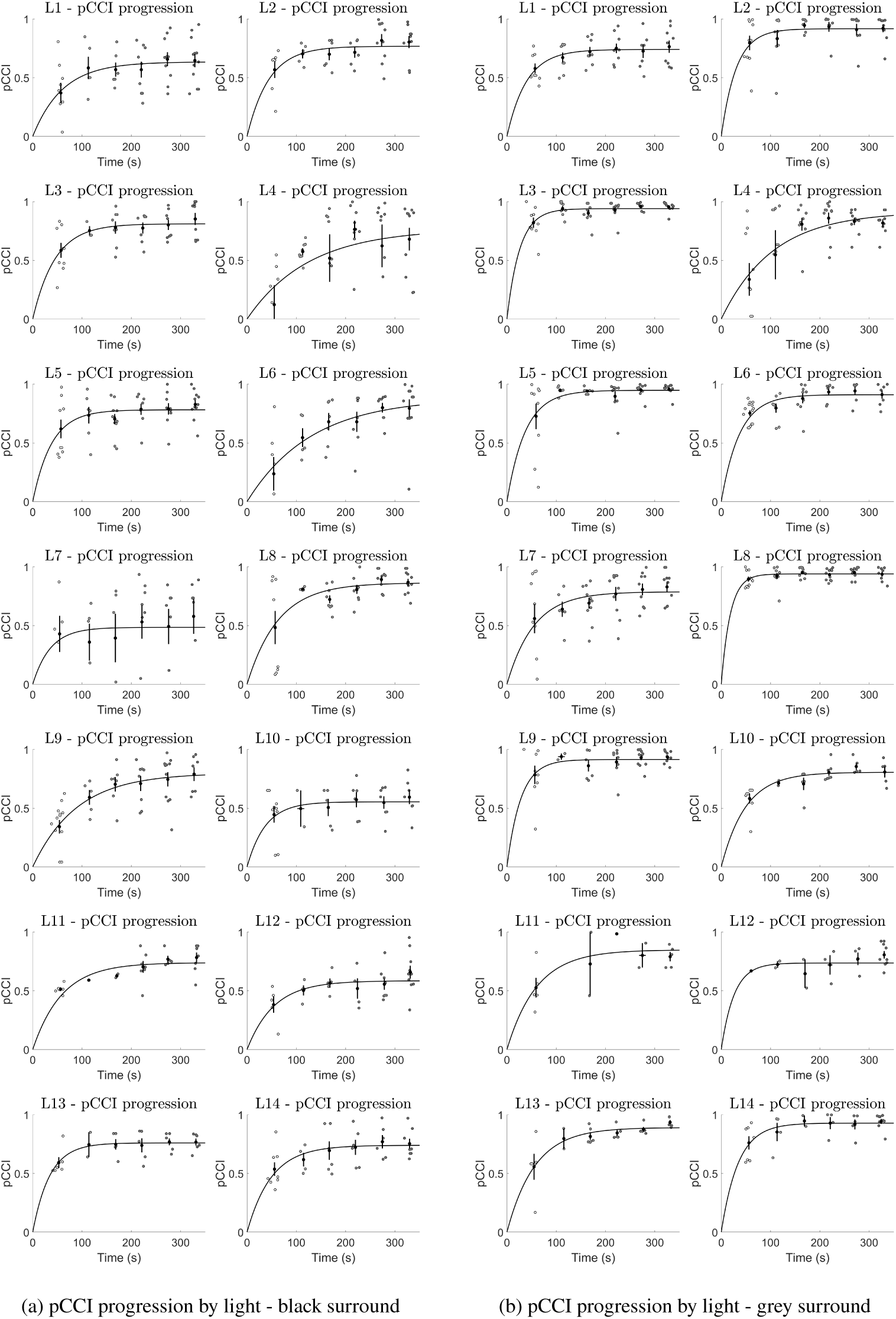
Submitted match pCCIs pooled over all participants, per light: (a) black surround, (b) grey surround. Grey and black markers indicate individual pCCI values and timepoint means, respectively. Error bars: standard errors of timepoint means.

**Fig. 13.**
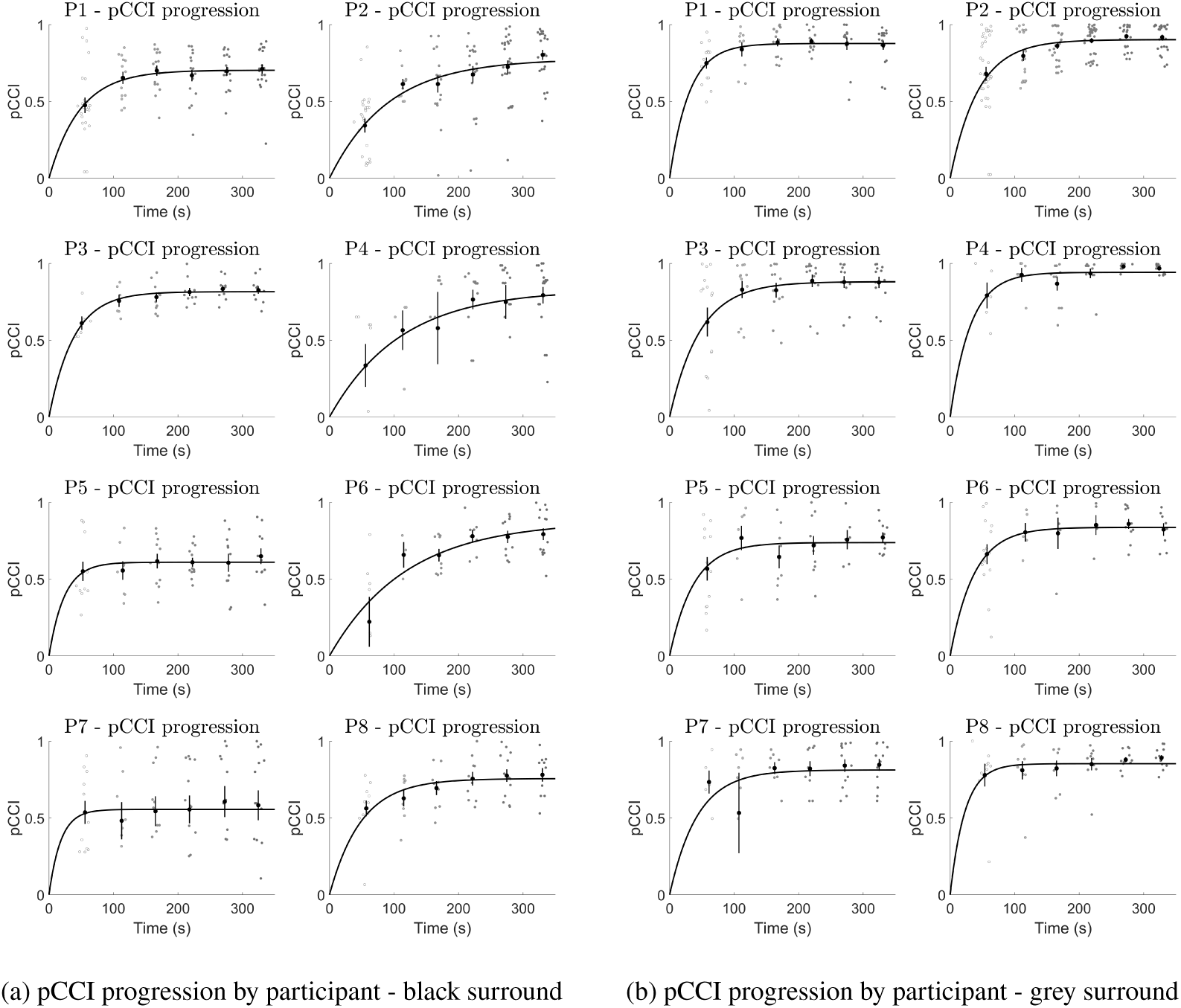
Submitted match pCCIs pooled over all lights, per participant: (a) black surround, (b) grey surround. Grey and black markers indicate individual pCCI values and timepoint means, respectively. Error bars: standard errors of timepoint means.

## Appendix C Illumination-final matches per light

**Fig. 14.**
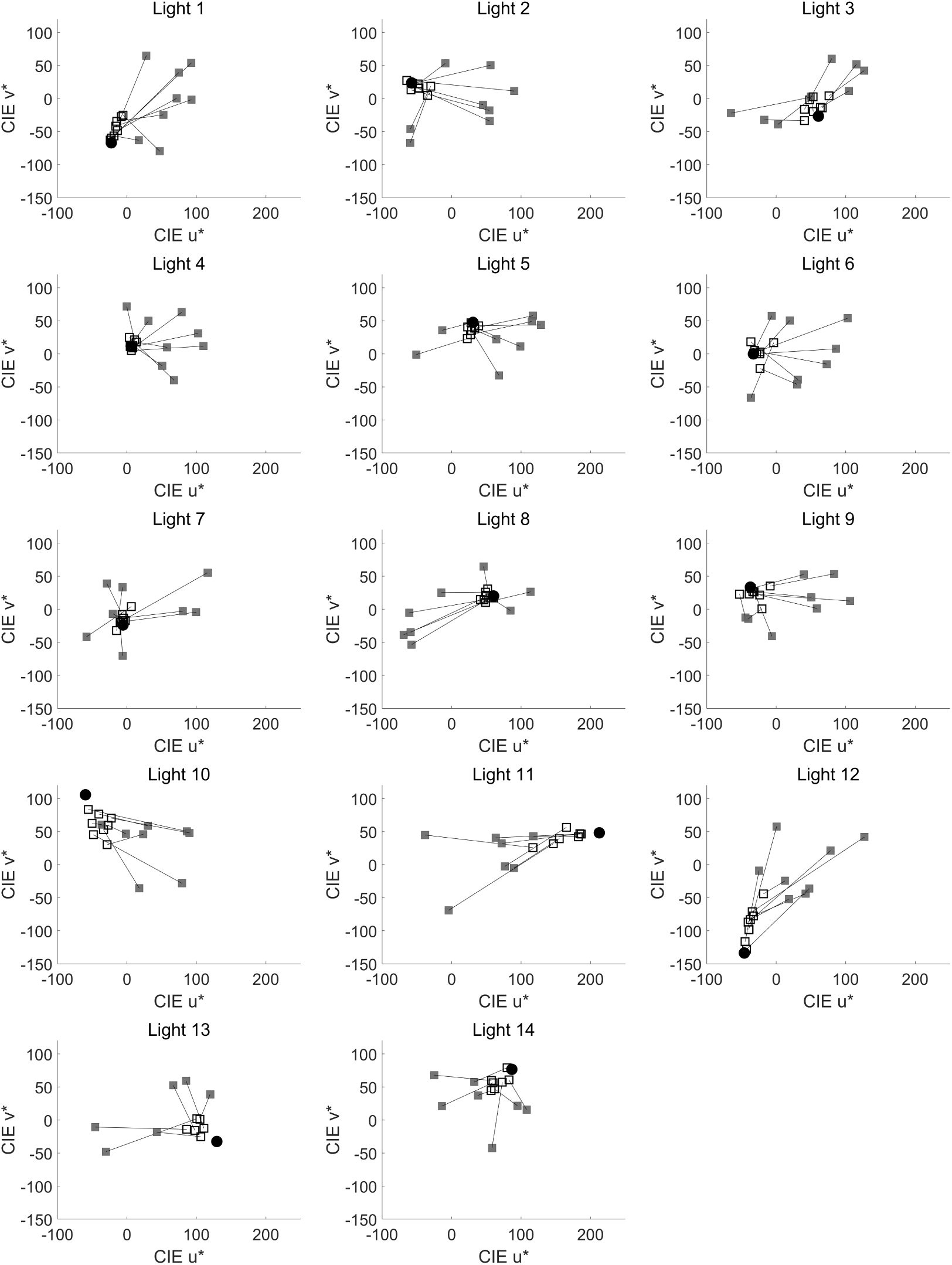
Illumination-final matches - black surround. Black circles are the test illumination chromaticity, grey solid squares the randomised start chromaticity for each trial, and black squares the illumination-final confirmed match for each trial. Lines join each pair of start and final chromaticities.

**Fig. 15.**
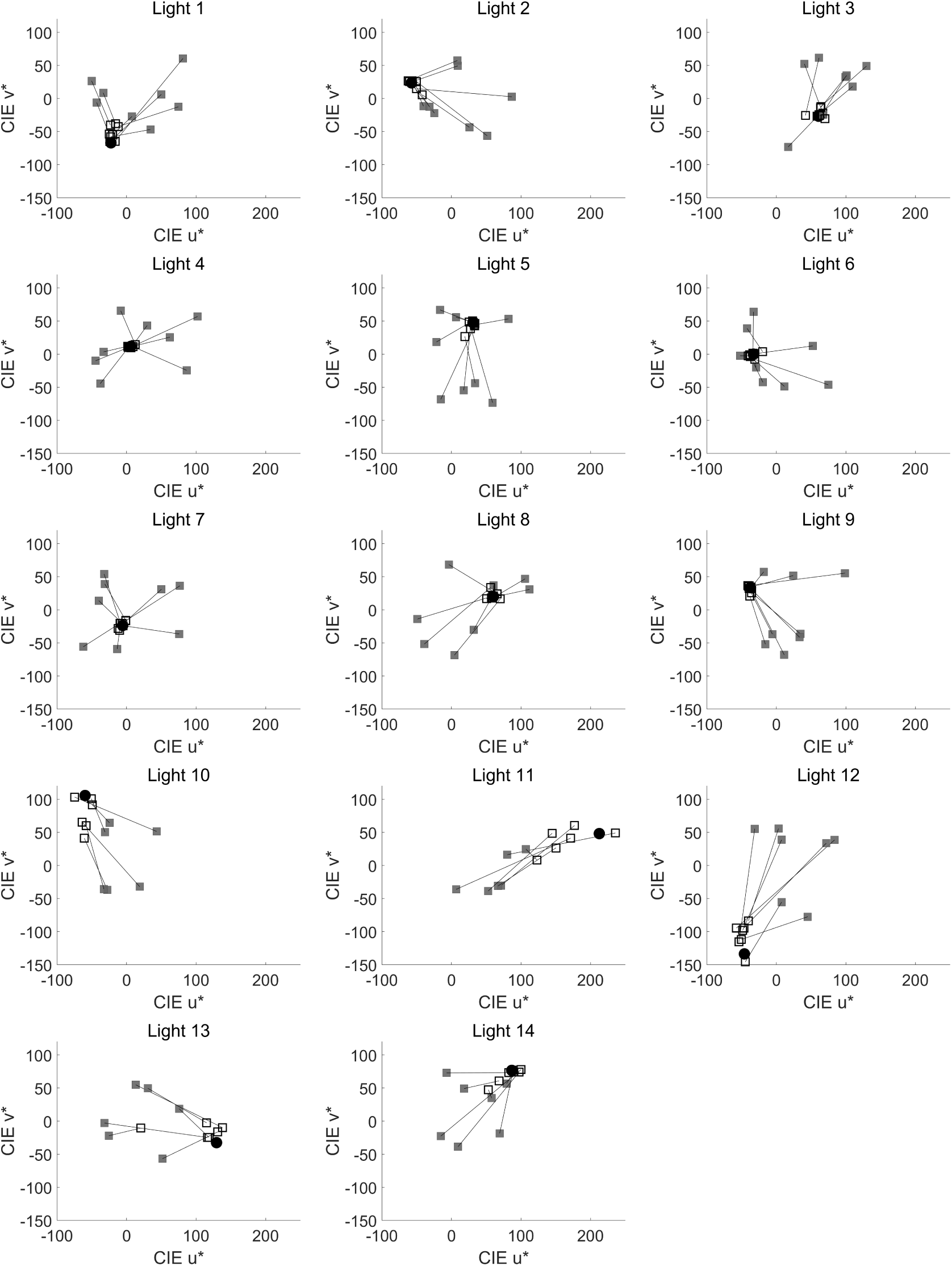
Illumination-final matches - grey surround. Black circles are the test illumination chromaticity, grey solid squares the randomised start chromaticity for each trial, and black squares the illumination-final confirmed match for each trial. Lines join each pair of start and final chromaticities.

## Footnotes

1 We will hereafter specify colorimetric differences using the form 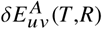, where *A* denotes the adapting chromaticity, and *T* and *R* are the two chromaticities being compared for colour difference.

2 We position the observer to have a perpendicular gaze to the surface of the viewing target, and calibrate the device screen at the same perpendicular angle of view. Radiance spectrometers are known to have increasing error, and electronic displays to produce shifted colours, with distinctive hue shifts by display type [32], at larger incident angles. The recommended standard practice is to calibrate perpendicular to the target surface. For experimental setups with non-perpendicular viewing angle, there is a trade-off between viewing position accuracy and radiance spectrophotometer accuracy.

3 The six moderate illuminations were chosen to coincide with discretely sampled chromaticities from Wiess et al. (2017) [23]

4 The blackbody and daylight loci are distinct but proximal for colour temperatures above 5000K.

5 It is not necessary that model parameters obtained by fitting to pooled data will be equivalent to the mean of model parameters obtained by fitting individually to all data subsets, however, agreement between these values is an indicator of model appropriateness.

